# Non-canonical *Drosophila* X chromosome dosage compensation and repressive topologically-associated domains

**DOI:** 10.1101/427443

**Authors:** Hangnoh Lee, Brian Oliver

## Abstract

**Background:** In animals with *XY* sex chromosomes, *X-*linked genes from a single *X* chromosome in males are imbalanced relative to autosomal genes. To minimize the impact of genic imbalance in male *Drosophila*, there is a dosage compensation complex (MSL), that equilibrates *X-*linked gene expression with the autosomes. There are other potential contributions to dosage compensation. Hemizygous autosomal genes located in repressive chromatin domains are often de-repressed. If this homolog-dependent repression occurs on the *X*, which has no pairing partner, then de-repression could contribute to male dosage compensation.

**Results:** We asked whether different chromatin states or topological associations correlate with *X* chromosome dosage compensation, especially in regions with little MSL occupancy. Our analyses demonstrated that male *X* chromosome genes that are located in repressive chromatin states are depleted of MSL occupancy, however they show dosage compensation. The genes in these repressive regions were also less sensitive to knockdown of MSL components.

**Conclusions:** Our results suggest that this non-canonical dosage compensation is due to the same trans-acting de-repression that occurs on autosomes. This mechanism would facilitate immediate compensation during the evolution of sex chromosomes from autosomes. This mechanism is similar to that of *C. elegans*, where enhanced recruitment of *X* chromosomes to the nuclear lamina dampens *X* chromosome expression as part of the dosage compensation response in *XX* individuals.

## Background

Genes come in pairs and large-scale deviation from this state is detrimental, most probably as a result of disrupted gene expression balance [1,2]. Sex chromosomes are a peculiar exception to this general rule. In *XY* systems, males have what amounts to a heterozygous deletion of an entire chromosome, bearing ~20% of the genes in the case of Drosophila, with no impact on fitness. In such systems, compensation often rectifies gene dose effects as a way to maintain gene balance [3–5].

In *Drosophila melanogaster*, a male-specific complex called the Male Specific Lethal (MSL) complex, plays a role in increasing expression of genes from the single *X* chromosome relative to autosomes. MSL and perhaps other unidentified sources of compensation ultimately achieve remarkably equalized levels of *X-*linked gene expression in males with one *X* and females with two *X*s, as well as balancing *X* expression with the autosomes [6,7]. This boosting of expression by MSL complex is primarily achieved via enhanced elongation of transcription [8], but there also is evidence that RNA polymerase II (PolII) binding is increased by 1.2 fold at male *X* chromosome promoters [9–11]. The complex includes MSL-1, MSL-2, and MSL-3 proteins, Maleless (MLE), and Males absent on the first (MOF) proteins, and two non-coding RNAs, *roX1* and *roX2 [12]*. MOF has a histone acetyltransferase activity and functions in enhanced elongation of *X* chromosome gene transcription by acetylating histone H4K16 (H4K16Ac) [8]. Binding of MSL complex to the male *X* chromosome occurs at Chromosome Entry Sites (CES), also referred to as High-Affinity Sites (HAS) [13,14]. The sites contain GA-rich sequences, called the MSL-recognition element (MRE) [14].

A related scheme for *X* chromosome dosage compensation occurs in *C. elegans*, where *XX* worms are hermaphrodites and *X0* worms are males. In *XO* males, yield of *X* chromosome gene products is increased using various mechanisms (e.g. increased PolII recruitment, mRNA stability, or translation rate) in both males and hermaphrodites [15–18]. However, solving the gene production difference between autosomes and *X* chromosomes in males results in over-expression in *XX* animals. To manage this increased activity, *XX* hermaphrodite *C. elegans* have a dosage compensation complex that represses gene expression from both *X* chromosomes [5,16]. The *C. elegans* Dosage Compensation Complex (DCC) targets the *X* chromosomes and spreads from recruitment sites on the *X* [19]. Recruitment of DCC on *X* chromosome is linked to increased mono-methylation of histone H4K20 (H4K20me1) [20,21], as well as depletion of histone modifications that mark active transcription, such as H4K16Ac [16,22,23], and H2A.Z variant histone [24]. These epigenetic changes accompany topological remodeling of the *X* chromosomes [25] and reduced PolII recruitment at *X-*linked promoters in hermaphrodites. [3,5,26]. This remodeling includes nuclear sub-localization of the *X* chromosomes to the lamina, which is generally repressive. Disruption of the anchoring between heterochromatin and nuclear lamina re-localized *X* chromosomes more centrally in the nucleus and results in partial de-repression of the *X-*linked genes [27]. The modulation of H4K16Ac in animals with a single *X* is a conserved characteristic between *D. melanogaster* and *C. elegans* [22] although the *XX* mechanisms differ [23].

Intriguingly, nuclear architecture-level de-repression also occurs in autosomal dosage compensation in *D. melanogaster*. Genes within repressive “topologically associated domains” (TADs), which include Lamina-associated domains (LADs) involved in dosage compensation in *C. elegans* hermaphrodites, show better autosomal dosage compensation in *Drosophila* hemizygotes [28]. The effect of autosomal deletions is de-repression of the non-deleted genes *in trans*, as well as a spreading of de-repression into flanking regions within the LAD. This suggests that these repressive domains are built based on additive or synergistic cooperation between gene homologs. This observation is of particular interest for two reasons. First, the necessity of two homologs for the repression is reminiscent of chromosomal pairing-dependent events, such as pairing-sensitive silencing [29,30], or transvection [31,32]. In transvection, the existence of homologous chromosome in proximity leads to enhancer action *in trans* or insulator bypass *in cis* [31]. As such, chromosomal pairing may provide a mechanistic basis of how autosomal deletions result in the de-repression of non-deleted genes [28]. The absence of a pairing partner for the single *X* in males might therefore be consequential. Second, the repression at the two-dose state, and de-repression at one-dose state, is analogous to *X* chromosome dosage compensation in *C. elegans*. This led us to ask if the de-repression of one-dose genes from repressive domains occurs on *D. melanogaster X* chromosomes. If so, this would contribute to dosage compensation in males.

## Results

### *X-*linked repressive TADs genes display low expression levels, but are dosage compensated in males

To determine the overall structure of chromatin domains on the *X*, we used results from three previous studies that divided genome into repressive vs. non-repressive chromatin domains/TADs and LADs vs. non-LADs. LAD and DamID (DNA adenine methyltransferase Identification) based chromatin occupancy information was from *Kc* cells [33,34]. TAD information was from Hi-C conformation capture from mixed sex embryos [35]. From the Hi-C study, “Null” TADs were characterized by general lack of chromatin marks, except for a weakly enriched binding of an insulator protein, Suppressor of Hairy-wing [SU(HW)]. The LAD and “Null” TADs correspond and largely overlap with “Black” domain DamID work. The “Black” domain has increased signals of Histone H1, Effete (EFF), Suppressor of Under-Replication (SUUR) and Lamin B protein binding. These repressive TADs are known to share various characteristics [35], and there are significant overlaps among the identified gene sets (**Figure 1A**). For example, 63% of genes that are in LADs are also in Black domains, and 78% of genes that are in Black domains are in Null domains. We collectively refer to these overlapping domains as “repressive TADs”.

**Figure 1.**
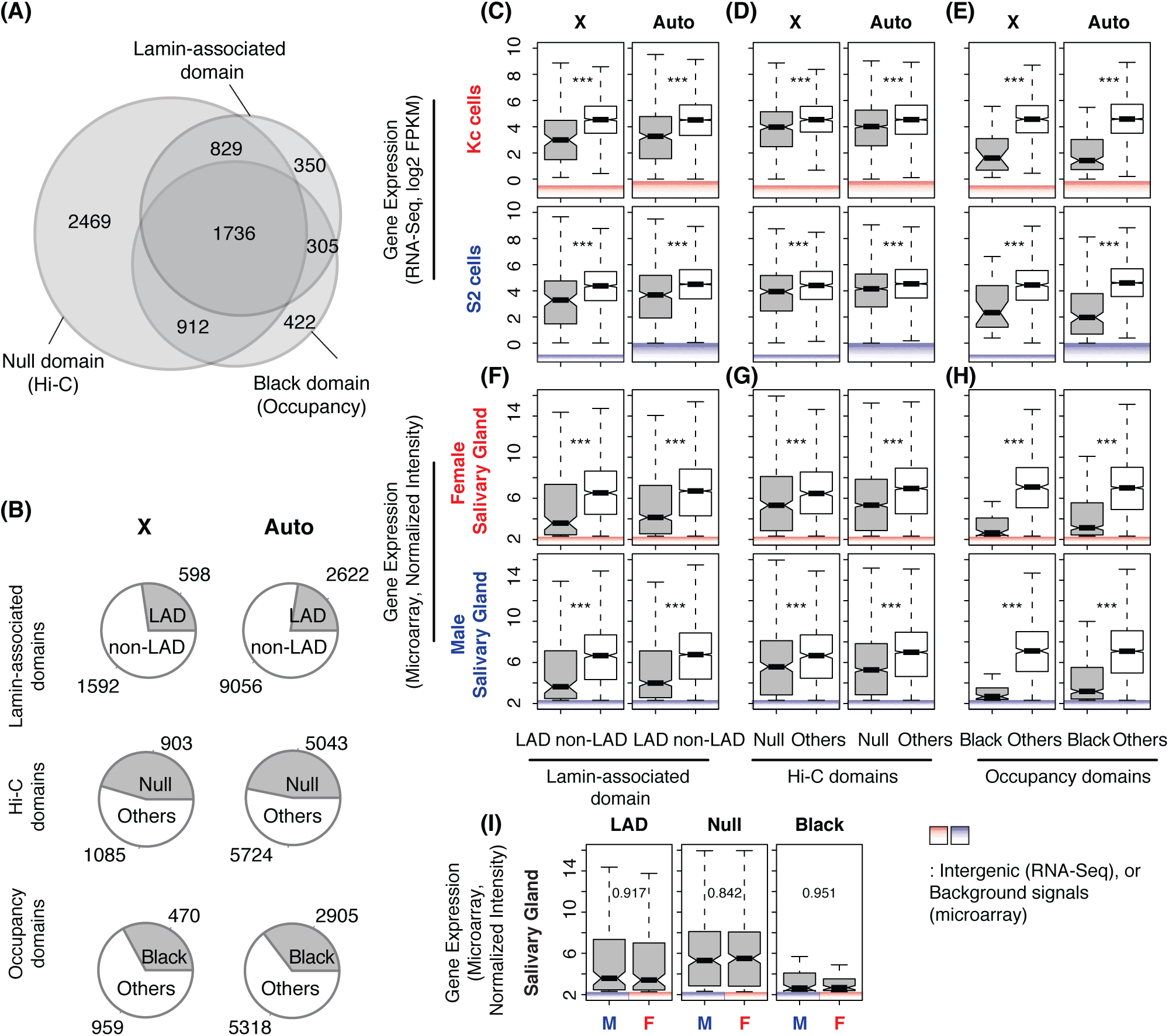
Repressive TAD genes display lower gene expression levels and are dosage-compensated in male cells. **(A)** Venn-diagram displays overlap among the three repressive TADs that are described in this study. **(B)** Pie charts demonstrate proportion of repressive TAD genes (gray) vs. non-repressive TAD genes (white) in *Drosophila* genome. In (A) and (B), only protein-coding polyA^+^ genes are counted. The numbers do not directly indicate numbers of “expressed” genes in each TAD. (**C-H**) Gene expression levels from the repressive TAD genes (gray) and non-repressive TAD genes (white) based on LAD (C,F), Hi-C (D,G), and chromatin occupancy studies (E,H). The top two rows show RNA-Seq results from *Drosophila* cell lines (unit: log2 FPKM, C-E), and the bottom two rows are from microarray study done with larval salivary glands (unit: normalized signal intensity, E-H). Intergenic signals from the 99th percentiles and below in RNA-Seq analyses, as well as background signals from the 95th percentiles and below in microarray results are indicated. **(I)** Comparisons of X chromosome gene expression levels from the repressive TADs between female and male salivary glands. Boxplots indicate the distribution of gene expression levels above expression cutoffs. Middle lines in box display medians of each distribution. Top of the box. 75th percentile. Bottom of the box. 25th percentile. Whiskers indicate the maximum, or minimum, observation within 1.5 times of the box height from the top, or the bottom of the box, respectively. Notches show 95% confidence interval for the medians. *** *p* < 0.001 from Mann-Whitney ⋃ test. The same format and test have been used for all boxplots appeared in this study.

Each of these three repressive TADs covered 23 to 47% of the protein-coding genes in the *Drosophila* genome. To describe which genes on the *X* were in repressive TADs, we parsed by chromosome (**Figure 1B**). Collectively, genes within LADs included 38% of *X* chromosome genes and 29% of autosomal genes (*p* = 1.19e-06, Fisher’s exact test). Genes within Null domains included 45% of *X* chromosome genes and 47% of autosomal genes (*p* = 0.25). Genes within Black domains included 33% of *X* chromosome genes and 35% of autosomal genes (*p* = 0.076). Clearly, a large fraction of the genome, including the X, are in repressive domains. If these genes are simply “off”, then asking if they are dosage compensated is a futile effort (2 × 0 = 0). Therefore, we carefully examined expression levels from genes that are within the repressive domains to see if we could reliably detect expression. We used previously reported expression data for this analysis [36,37]. Expression levels in repressive domains were reassuringly lower than in non-repressive domains. We found these trends of lower expression in repressive-TAD genes when we investigated different cell lines (**Figure 1C-E**) and sexed salivary glands (**Figure 1F-H**), but there was clear evidence of expression.

Determining the difference between low and off is critical for this analysis. We measured the biological and technical noise levels by measuring intergenic signals (**Figure 1C-E**). The 99th percentiles for intergenic signal were 0.87 Fragments Per Kilobase of transcript per Million mapped reads (FPKM) in *S2* cells (male) and 0.98 FPKM in *Kc* cells (female). This is in stark contrast to expression levels in the repressive TADs from *Kc* cells, where LAD and Black domains were determined (**Figure 1C-E**, the top panel). The median gene expression level was 9.1 FPKM for genes within LADs and 16.1 FPKM for genes within Null domains. Genes in Black domains showed lower expression at a median of 2.8 FPKM, but all of these expression levels far exceed our estimates of noise. In *Kc* cells, approximately 19.6% and 40.4% of the *X-*linked genes demonstrate gene expression above the cutoff levels from LAD and Null domains, respectively. Only 5.7% of the *X-*linked genes were expressed from Black domains, indicating that the Black domain has the most repressive characteristics among three different calls of repressive TADs. Autosomal genes from repressive TADs also displayed lower gene expression levels compared to non-repressive TAD genes with 9.7 (LAD), 16.1 (Null), and 2.7 FPKMs (Black), which are not significantly different from repressive TAD genes on the *X* (*p* > 0.2, Mann-Whitney ⋃ test). In male *S2* cells, the repressive TAD genes demonstrated 9.8 (LAD), 15.4 (Null), and 5.1 FPKMs (Black), underscoring that repressive TAD genes on the *X* have comparable expression levels between *S2* and *Kc* cells (*p* > 0.05, **Figure 1C-E**), and are thus dosage compensated. We made a similar observation from sexed salivary glands. A large fraction of genes from repressive TADs showed higher expression than the technical noise level, which we determined based on backgrounds signals from the control probes of microarrays (**Figure 1F-H**, normalized intensities of approximately 2.3 in both sexes). For example, about 27.4% and 26.6% of the total *X-*linked LAD genes showed gene expression above the background levels in female and male salivary glands, respectively. Thus, we were confident that we could make meaningful measurements of dosage compensation among those genes. Briefly, a substantial portion of the genes in repressive TAD domains showed detectable levels of gene expression. We used these genes in our analysis.

Genes in repressive TADs demonstrated comparable expression levels between female (*Kc*) and male (S2) cells from the *X* (**Figure 1C-E**), indicating that they are dosage compensated. To confirm that male *X-*linked genes are dosage compensated in repressive TADs, we compared expression profiles from salivary glands, where female and male were matched siblings. From microarray results, we observed that male *X-*linked genes from LAD regions demonstrated comparable expression levels to that of females (**Figure 1I**). The median signal intensity from male *X-*linked genes was 3.41, which did not differ from that of female (3.58, *p* = 0.917) despite the 50% difference in *X* gene dose. We obtained similar equilibrated expression of the *X* from other repressive TADs. *X* chromosome genes in the Null domains showed a median of 5.51 signal intensity in *X* males when it was 5.31 in *XX* females. Black domain genes had medians of 2.68 and 2.64 signal intensities in *X* males and *XX* females, respectively. Overall gene expression signals from autosomes were consistent between two sexes (6.52, *p* > 0.996 for differential expression). Therefore, the repressive TAD genes are dosage compensated in males.

### Repressive TAD genes lack MSL complex binding

*X* specific dosage compensation has canonical and non-canonical components. If canonical dosage compensation is active in repressive domains, the MSL complex should occupy those regions. To address this possibility, we first investigated chromatin occupancy by MOF, the key writer of the H4K16Ac mark [38] in the MSL complex [12]. MOF also has an MSL-independent role in regulating a smaller subset of genes in both sexes by participating in Non-Specific Lethal (NSL) complex [39]. We analyzed genome-wide Chromatin Immunopreciptation (ChIP) results [36,40] to determine occupancy of the MSL complex as well as H4K16Ac levels in tissue culture cells and salivary glands (we measured MOF and H4K16Ac enrichment within gene bodies because both MOF and H4K16Ac display broad enrichment patterns over these features [39]). Strikingly, in male *S2* cells, MOF binding in *X* chromosome repressive TADs was significantly lower than elsewhere on the *X* (*p* < 6.01e-4) and was comparable to occupancy levels on the *X* in female *Kc* cells (**Figure 2A-C**). H4K16Ac enrichment concurred with MOF occupancy. In all classes of *X* chromosome repressive TADs, H4K16Ac levels were significantly lower than in other domains (*p* < 6.96e-09, **Figure 2D-F**). H4K16Ac levels on *X-*linked genes was still higher in *S2* cells than *Kc* cells even within repressive TADs (*p* < 3.02e-09). However, the differences in H4K16Ac levels between male and female cells were significantly smaller in LAD and Null domains than non-repressive TADs on the *X* (*p* < 7.14e-04, Permutation test). The Black domain genes showed the same trend, although this was not statistically significant (*p* = 0.27). Consistent with the MOF occupancy and H4K16Ac enrichment, we observed that MSL-1 occupancy was lower in genes within repressive TADs on the *X* compared to non-repressive TADs (*p* < 1.11e-12, Mann-Whitney ⋃ test, **Figure 2G-I**). Thus, the occupancy and activity of MSL complex is reduced in the case of the dosage compensated *X-*linked genes in *S2* cell repressive TADs.

**Figure 2.**
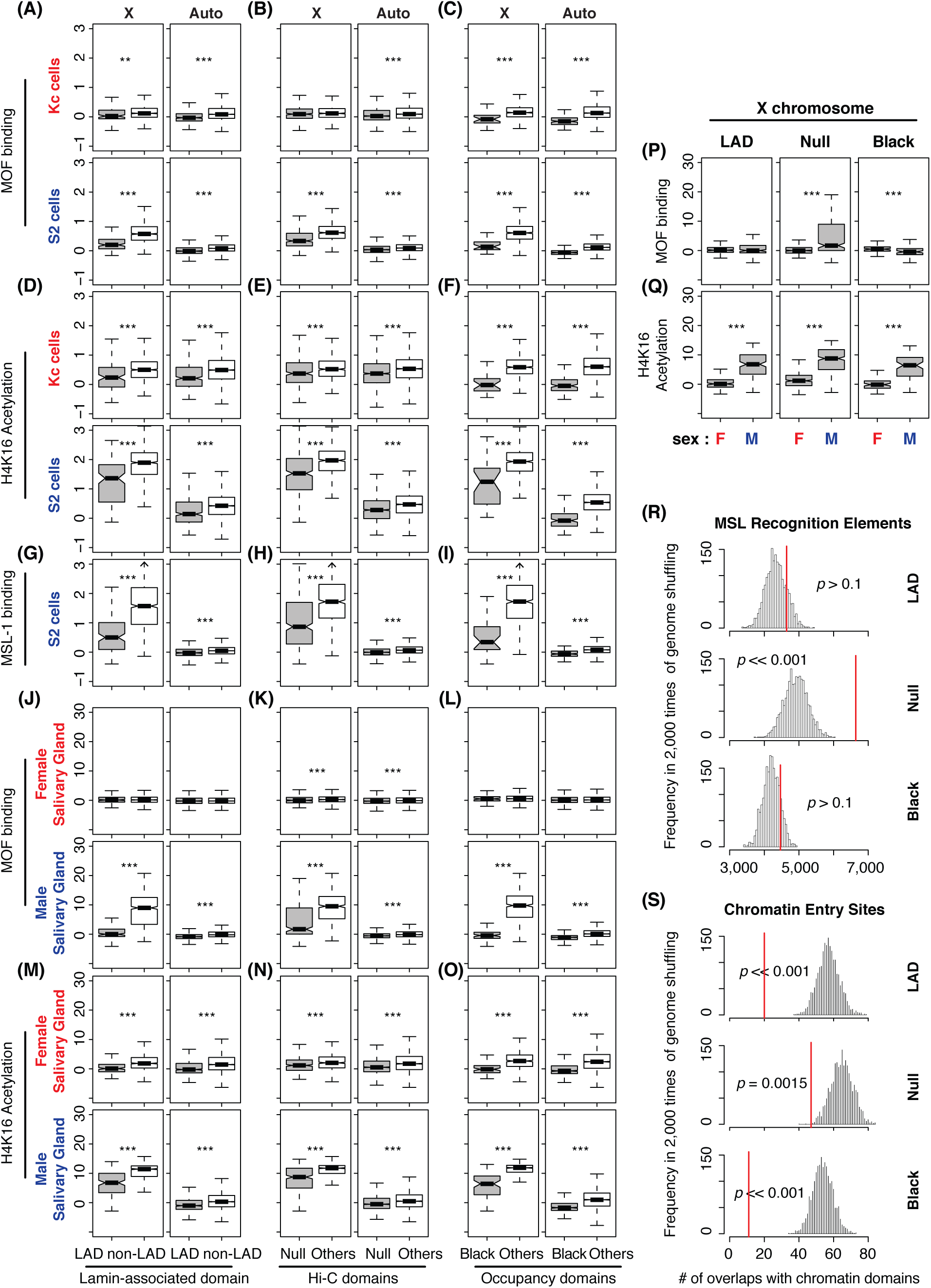
Repressive TAD genes have a limited binding of MSL complex. **(A-I)** Chromatin immunoprecipitation (ChIP) results from MOF binding (A-C), Histone H4K16 acetylation (D-F), and MSL-1 binding (G-I) are summarized as boxplots for *Drosophila* cell lines (*Kc* and *S2*). Gene level ChIP signals are separately shown based on LAD (A,D,G), Hi-C (B,E,H) and chromatin occupancy (C,F,I) study results. **(J-O)** ChIP results from the 3rd instar larval salivary glands. ** *p* < 0.01, *** *p* < 0.001. **(P, Q)** Direct comparisons of MOF binding (P) and H4K16Ac enrichment (Q) between female and male salivary glands from (J-O). (**R, S**) The histogram represents expected numbers overlaps between repressive TADs and MRE (R), or CES (S). We performed random shuffling of the X chromosome genome 2,000 times and demonstrated the frequencies of the numbers of overlaps. Red lines. the actual number of overlaps between LADs, and MREs or CES’s. *p* values are from permutation tests.

To examine MSL complex activity at the repressive TADs in tissues, we analyzed ChIP results from sexed larval salivary glands. In males, X chromosome MOF binding was significantly higher at gene bodies in non-repressive TADs, compared to repressive TADs (**Figure 2J-L**, *p* < 2.2e-16). If MOF binding is functional, then the H4K16Ac mark should follow a matching enrichment pattern. Indeed, H4K16Ac levels were higher at genes in non-repressive TADs compared to repressive TADs (**Figure 2M-O**, *p* < 2.2e-16). As we observed in the tissue culture cells, the basal level of H4K16Ac was higher in repressive TADs of the male salivary glands, compared to that of female glands, despite the fact that MOF occupancy was not significantly higher in LAD and Black domains (*p* > 0.63, **Figure 2P,Q**). This observation indicates that regulation of repressive TAD genes on the *X* chromosome occurs with limited or transient access to MSL complex, but can involve modulations of H4K16Ac in a canonical manner. We explore the degree of function associated with this lower H4K16ac in a later section.

Since the genes within repressive TADs have low occupancy of MSL complex and lower H4K16ac, we wondered if repressive TADs lack genomic signatures that are required for MSL complex binding. Specifically, we asked if lower MOF activity correlates with lower density of the MSL complex entry sites in repressive domains. *Drosophila* MSL complex specifically binds to X, which occurs at CES [14]. CES contains GA-rich DNA sequence motif, called MRE, whose introduction to an autosome results in local recruitment of MSL complex to that site [14]. We identified 11,306 MRE motifs from the *X* chromosome of the reference genome (using an *E*-value < 10e-5 cutoff). The number of *X* chromosome MREs in repressive domains was not statistically different from random (**Figure 2R**, *p* > 0.1 Permutation test), indicating that the repressive TADs are not free of MRE motifs. However, when we investigated if genes in repressive TADs recruit MSL complex to their chromatin regions, we found only 20 overlaps between LADs and the 150 CES (approximately 57 expected, *p* << 0.001, Permutation test, **Figure 2S**) that recruit MSL [14]. We obtained consistent results from Null and Black domains (**Figure 2R,S**). These observations suggest that, on male *X* chromosomes, MSL complex does not efficiently bind genes within the repressive TADs.

### X-linked repressive TADs genes are less sensitive to disruption of MSL-complex functions compared to the canonical dosage-compensation target genes

If the repressive TAD genes are dosage-compensated in a non-canonical way on the male *X* chromosome, such genes might be indifferent to MSL complex function. If the low level of H4K16ac is matched to the generally low level expression of the genes in repressive domains, compensation of such genes should depend on MSL function. To investigate the impact of disrupted MSL complex function on *X-*linked genes in repressive TAD domains, we analyzed gene expression profiles of S2 cells whose MSL components were selectively depleted via RNAi-mediated knockdown [36,41,42]. When *mof* mRNA was depleted, *X-*linked genes within LADs displayed significantly less gene expression reduction; approximately 1.2 fold higher than genes in non-LAD domains (*p* = 1.1e-13, Mann-Whitney ⋃ test, **Figure 3A**). We made similar observations from *X* chromosome genes that belong to Null and Black domains from the Hi-C study and occupancy study. They exhibited about 1.1 to 1.3 fold more expression upon the depletion of *mof* than other *X-*linked genes in non-repressive domains (*p* < 2.3e-08). As expected, those chromatin regions that lack MOF binding and H4K16Ac were less sensitive to the RNAi treatment as well. Genes from regions without enrichment of MOF or H4K16Ac showed 1.2 fold more expression than the enriched regions (*p* = 1.1e-14 for MOF, and 0.11 for the acetylation). MOF is also bound to sites on autosomes as a part of NSL complex, while it activates only a small subset of genes that the complex binds to [43]. Consistent with this idea, we saw little down-regulation of overall autosomal gene expression from the *mof* depleted *S2* cells (*p* > 0.05, **Figure 3B**).

**Figure 3.**
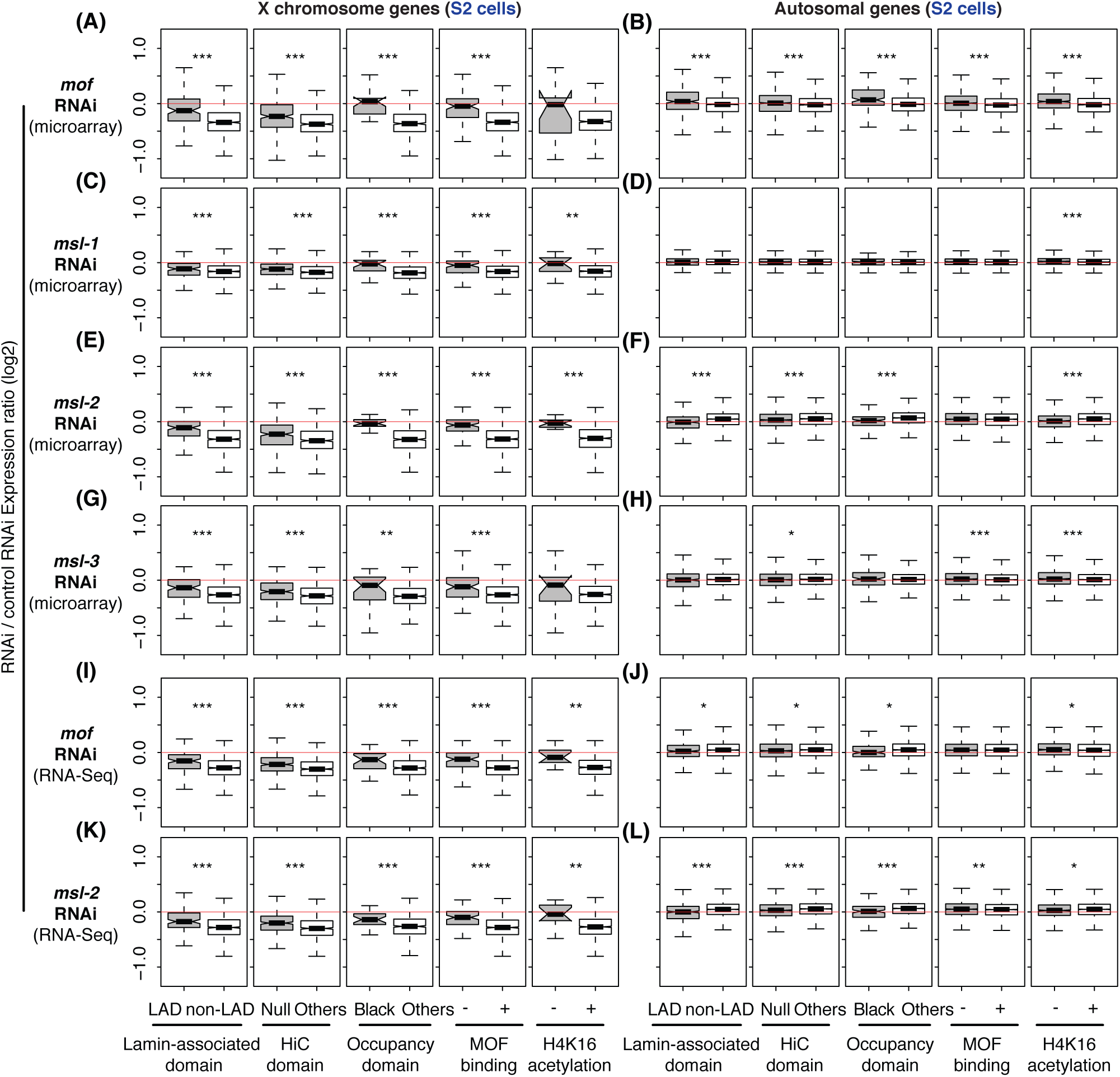
Different responses from the repressive vs. non-repressive TAD genes upon knockdown of a MSL complex component. Boxplots represent gene expression changes in log2 scale from depletion of MSL components in *Drosophila S2* cells. Plots are based on three independent studies [36,41,42], which used either microarray (A-H) or RNA-Seq technology (I-L). **(A, B)** Differential gene expression from *mof* knockdown cells. Changes from the repressive TADs (left three columns, LAD, Null, and Black) as well as MOF binding, or Histone H4K16 acetylation regions are presented. **(C-H)** Results from *msl-1, msl-2*, or *msl-3* knockdown. **(I, J)** Results from *mof* knockdown, measured by RNA-Seq analysis. **(K, L)** *msl-2* knockdown. **(A,C,E,G,I,K)** Changes from *X* chromosome genes. **(B,D,F,H,J,L)** Changes from autosomal genes. * *p* < 0.05, ** *p* < 0.01, *** *p* < 0.001

We also asked if the expression of *X-*linked genes in repressive TADs was less sensitive to depletion of other MSL components. Our analysis showed significantly less reduction in expression in gene within repressive TADs, relative to non-repressive TADs, when *msl-1* mRNA was depleted (**Figure 3C,D**, *p* < 0.001). Similarly, *msl-2* and *msl-3* knockdown caused 1.1 to 1.2 fold more *X* chromosome gene expression from genes in repressive TADs, compared to non-repressive TADs (**Figure 3E-H**, *p* < 0.01). These results were not due to the inaccurate detection of low abundant transcripts in hybridization-based techniques (i.e. microarrays) [44,45]. When we analyzed an independent study that performed RNA-Seq analysis of either *mof* or *msl-2* depleted *S2* cells, we also observed about 1.1 fold more expression from the *X-*linked genes within repressive TADs compared to non-repressive TADs (*p* < 0.001, **Figure 3I-L**). Therefore, results from the MSL knockdown were consistent with our observation in **Figure 2** and suggest that repressive TAD genes on the *X* chromosome do not rely entirely on MSL complex for dosage compensation.

We inspected MOF occupancy and H4K16Ac enrichment at individual genes in repressive TADs on the *X* to determine the patterns of occupancy across the gene bodies. Genes that were clearly regulated by the canonical dosage compensation machinery (i.e. MSL-dependent) display broad enrichment signals of MOF and H4K16Ac across the gene body regions, whereas MSL-independent MOF target genes (e.g. MOFs in NSL complex) show promoter-enriched MOF binding patterns [39]. We observed genes that were sensitive to the *msl* or *mof* knockdown, for example *CG9947* and *arm*, had broad ChIP signals of MOF and H4K16Ac in contrast to an autosomal gene, *RpL32*, which has MOF enrichment only at its promoter region (**Figure 4A-C**). Compared to those canonical MSL target genes, the genes in repressive TAD regions showed absent MOF binding (**Figure 4D,E**, *CG34330* and *CG9521*), or weak MOF occupancy (**Figure 4F,G**, *CG8675* and *CG2875*). In all four specific cases, the knockdown of *mof* or *msl-2* did not lead to statistically significant reduction of gene expression in males (*p* > 0.7, **Figure 4D-G**), additionally the genes were still fully compensated relative to females in the salivary glands [male/female expression ratios of 1.02 (*CG34330*), 1.02 (*CG9521*), 1.04 (*CG8675*), and 0.97 (CG2875)]. For the latter class of genes that have weak MOF occupancy (*CG8675* and *CG2875*), we noticed that MOF also bound at the 3’ ends of genes. Furthermore, H4K16Ac enrichment had peaks at the 3’ ends as well, which contrasted with the promoter-focused peaks from NSL-MOF target genes (*RpL32*, **Figure 4A**), suggesting that there was some residual MSL activity, rather than NSL. The overall insensitivity to MSL RNAi-depletion suggests that the dosage compensation of these genes does not rely on the MSL complex, but requires additional mechanisms, such as de-repression.

**Figure 4.**
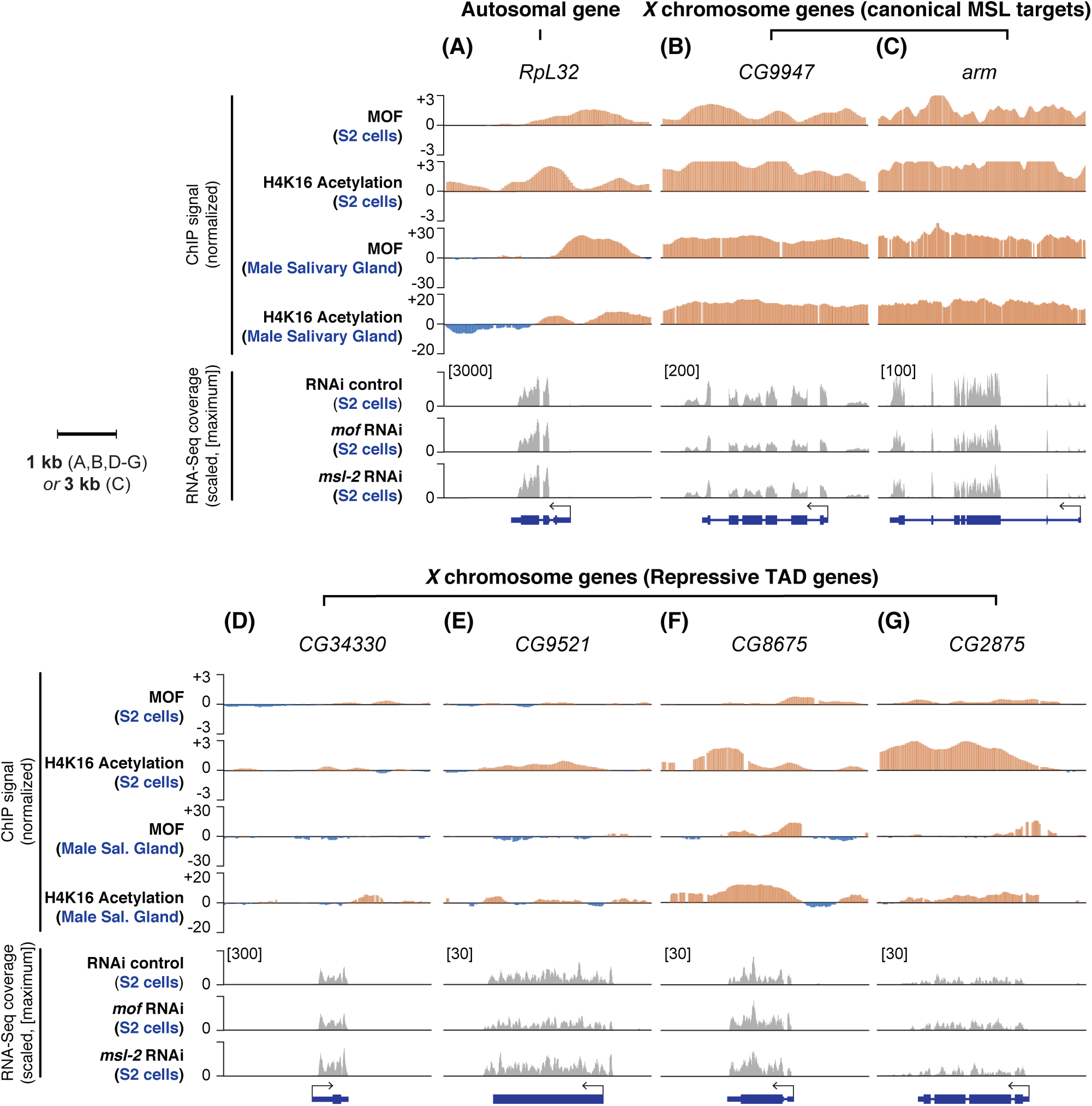
Repressive TAD genes that are less sensitive to *msl* or *mof* knockdown lack MOF enrichment at their gene bodies. Top four panels demonstrate normalized ChIP signals of MOF and H4K16Ac from *S2* cells as well as male salivary glands. The bottom three panels display RNA-Seq read coverages from our re-analysis of Zhang et al [42]. The plots are scaled based on their maximum coverage from one of the three samples: control RNAi, *mof* RNAi, and *msl-2* RNAi (indicated in the square brackets). Note that there is no sample-to-sample normalization because the total number of reads are similar across the three samples; 7.2, 7.5 and 8.1 million mapped reads for the control, *mof* RNAi, and *msl-2* RNAi samples, respectively. **(A)** An autosomal gene (*RpL32*). **(B, C)** Canonical MSL targets genes. *CG9947* (B) and *arm* (C). **(D-E)** Repressive target genes that are compensated via the non-canonical dosage compensation. *CG34430* (D), *CG9521* (E), *CG8675* (F), and *CG2875* (G).

### Non-canonical dosage compensation is more evident within TAD boundaries that are maintained between male and female cells

TAD boundaries are stable across different cell types, and even display evolutionary conservation [46]. Hi-C studies from *Kc* and *S2* cells showed that approximately 74% of TADs are located at the identical positions between the two cell lines [47]. The overall organization of *X* chromosome TADs is highly similar between the two cell lines, and depletion of *msl-2* or *msl-3* does not alter chromatin conformation of the *X* [48]. Nevertheless, it is still possible that the compensation of repressive TAD genes, as well as the increase of histone H4K16Ac (**Figure 2D-F, M-O**), are due to topological differences of chromatin between the two cell lines because the repressive TADs are originally defined from *Kc* cells as well as mixed sex embryos. Testing the possibility is important in this study for two reasons. First, it is possible that our observation could result from erroneous mapping of *Kc* cell-based TAD calls to *S2* cells, although this seems unlikely as MOF occupancy signal is very weak over the repressive TADs in the both cell lines (**Figure 2A-C, J-L**). Second, considering that CESs are enriched at TAD boundaries, from where the MSL complex spreads out based on proximity [48], if TAD structures differ between male and female cells, the architectural difference could contribute to the non-canonical dosage compensation.

To test the effects of potentially different TAD structure between lines and/or sexes, we investigated *X-*linked gene dosage compensation where TAD locations did not match between the two cell lines. For this purpose, we subclassified repressive TADs into four groups [47], which divided regions that are: within TAD boundaries in both *Kc* and *S2* cell lines (TT); either at boundaries or interspace between two large TADs in both cell lines (II); and within large TADs from only one of the cell lines (TI or IT, **Figure 5A**). The proportion of mismatches in TADs and their boundary locations (e.g. TT and II versus TI and IT) did not show bias based on autosome and *X* chromosome location, even in repressive TAD regions (*p* > 0.1 for all repressive TADs, Fisher’s exact test with Bonferroni correction, **Figure 5B-D**). This observation indicates that sex-specific alteration of TAD structures are unlikely to drive non-canonical male *X* chromosome dosage compensation. We also investigated the possible function of sex-specific TADs, by asking if knockdown of MSL components had a greate effect on repressive TADs domains specific to male *S2* cells (IT). However, this was not the case. Instead, we observed that the genes within repressive TAD domains and boundaries whose TAD locations were well-matched between the two cell lines (TT and II) were better compensated than the others (TI and IT) after depleting MSL components (**Figure 5E-G**). For example, LAD-associated genes from TT or II class regions demonstrated excellent dosage compensation after *msl-2* knockdown compared to the same genes from the control RNAi samples (0.94 fold, *p* = 0.047, Mann-Whitney ⋃ test, **Figure 5E**). We observed the same trend when we looked at genes from Null and Black domains (**Figure 5F,G**). In conclusion, these results suggest that our observation of non-canonical dosage compensation is not due to sex-specific modification of large TAD structures between male and female cells. TAD that differ between female *Kc* and male *S2* cells may be due to the rearrangements in these highly aneuploid cells, not differences in sexual identity.

**Figure 5.**
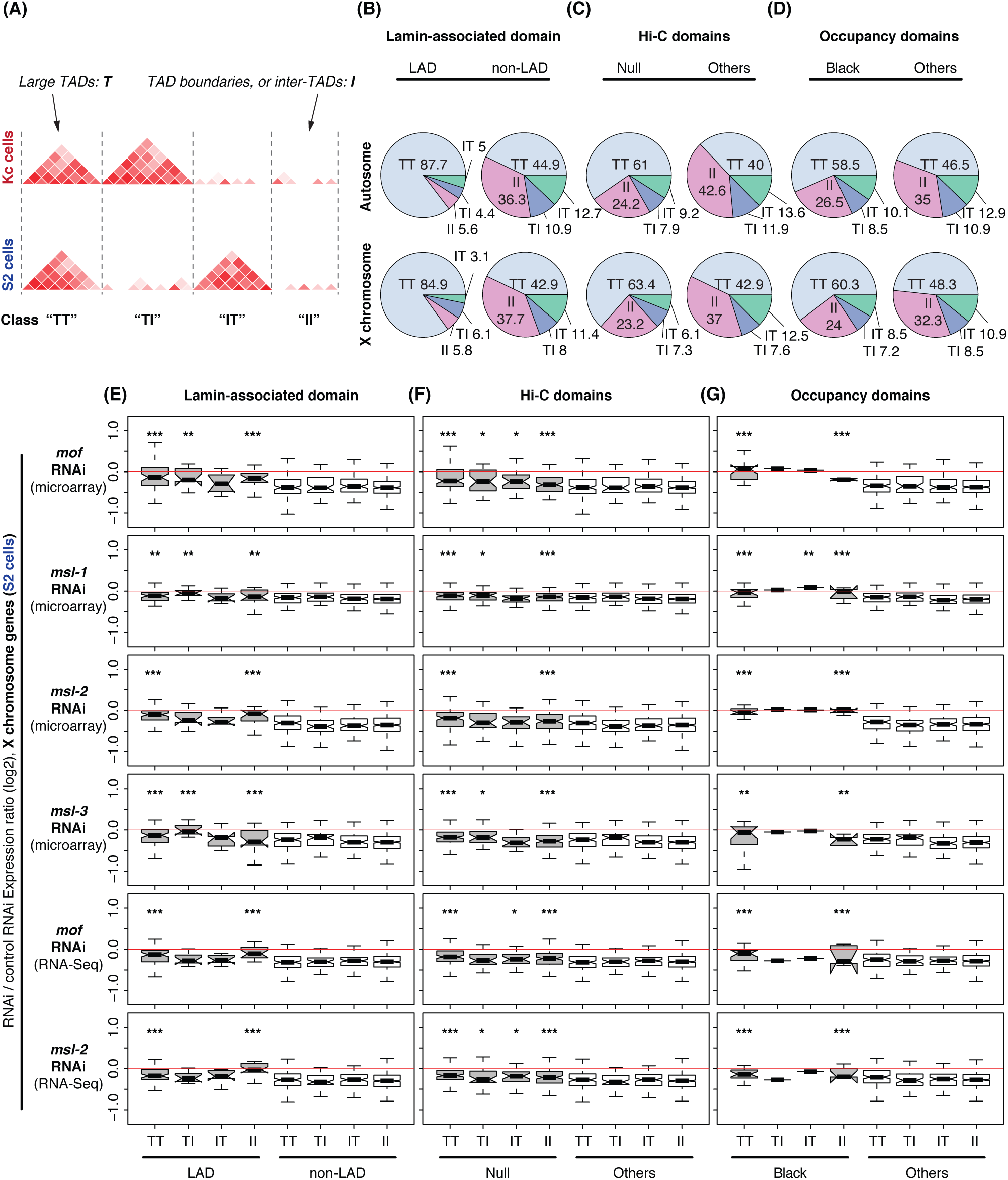
Repressive TAD-based dosage compensation occurs at where X chromosome TAD structures are maintained between female and male cells. (A) Schematic illustration of classification of TAD differences between *Kc* and *S2* cells based on [47] (modified with permission by the authors) **(B-D)** Percentage of the number of 2kb windows from four different classes of TAD difference between *Kc* and *S2* cells (TT, II, TI, and IT); windows that are found from within TAD boundaries in both *Kc* and *S2* cells (TT), that are at boundaries or interspace between TADs in both *Kc* and *S2* cells (II), that are found within the TAD boundaries in *Kc* cells but not in *S2* cells (TI), and that are not in TAD boundaries in Kc cells but within TAD boundaries in *S2* cells (IT). Top. distribution of 2kb windows from all autosomes. Bottom. Distribution of 2kb windows from X chromosomes. **(E-G)** Gene expression changes of *X-*linked genes in *S2* cells upon RNAi knockdown of *mof, msl-1, msl-2*, and *msl-3* as appeared in Figure 3 but based on TAD difference classes (TT, II, TI, and IT). Changes from repressive TADs (grey) and non-repressive TADs (white) are displayed. *P* values indicate differences from the median gene expression fold changes of non-LAD associated genes. * *p* < 0.05, ** *p* < 0.01, *** *p* < 0.001

## Discussion

A subset of *X* chromosome genes that are unbound by MOF still dosage compensate [49]. We have studied *X* chromosome dosage compensation of genes within repressive TADs in *Drosophila*, and their association with MSL dosage compensation complexes and activities. Our results revealed that genes from such repressive TADs are compensated with minimal contributions from MSL. We suggest those regions are able to achieve dosage compensation due to the weaker repressive TADs on the unpaired male chromosome.

This non-canonical compensation may be the same as observed in the case of autosomal deletions within or at the boundaries of repressive TADs [28]. These deletions have a dominant de-repressing effect, which results in partial dosage compensation for the hemizygous segment, and over-expression of genes in flanking two-dose regions (**Figure 6A,B**). These data suggest that repressive domains are established, strengthened, or stabilized by the existence of homologous pairs of chromosomes. There is strong precedent for pairing-dependent mechanisms in *D. melanogaster* that are known to activate or repress genes when homologous chromosomes are proximally located [29–32]. We suggest that the unpaired *X* chromosomes of males have weaker repressive domains than the same domains in the paired *X* chromosomes of females (**Figure 6C,D**). Thus, one can think of this as dosage compensation mediated by partial *X* inactivation in females, with de-repression in males. This model hinges on the reorganization of the nuclear lamina-DNA interaction, which can clearly regulate gene activities during cell differentiation even in the absence of global changes of the nuclear architecture [50]. For example, in mouse embryonic stem cells, loss of the tethering in the *Hdac3* deletion releases genomic regions of lineage-specific genes from nuclear lamina resulting in precocious expression of those genes [51].

**Figure 6.**
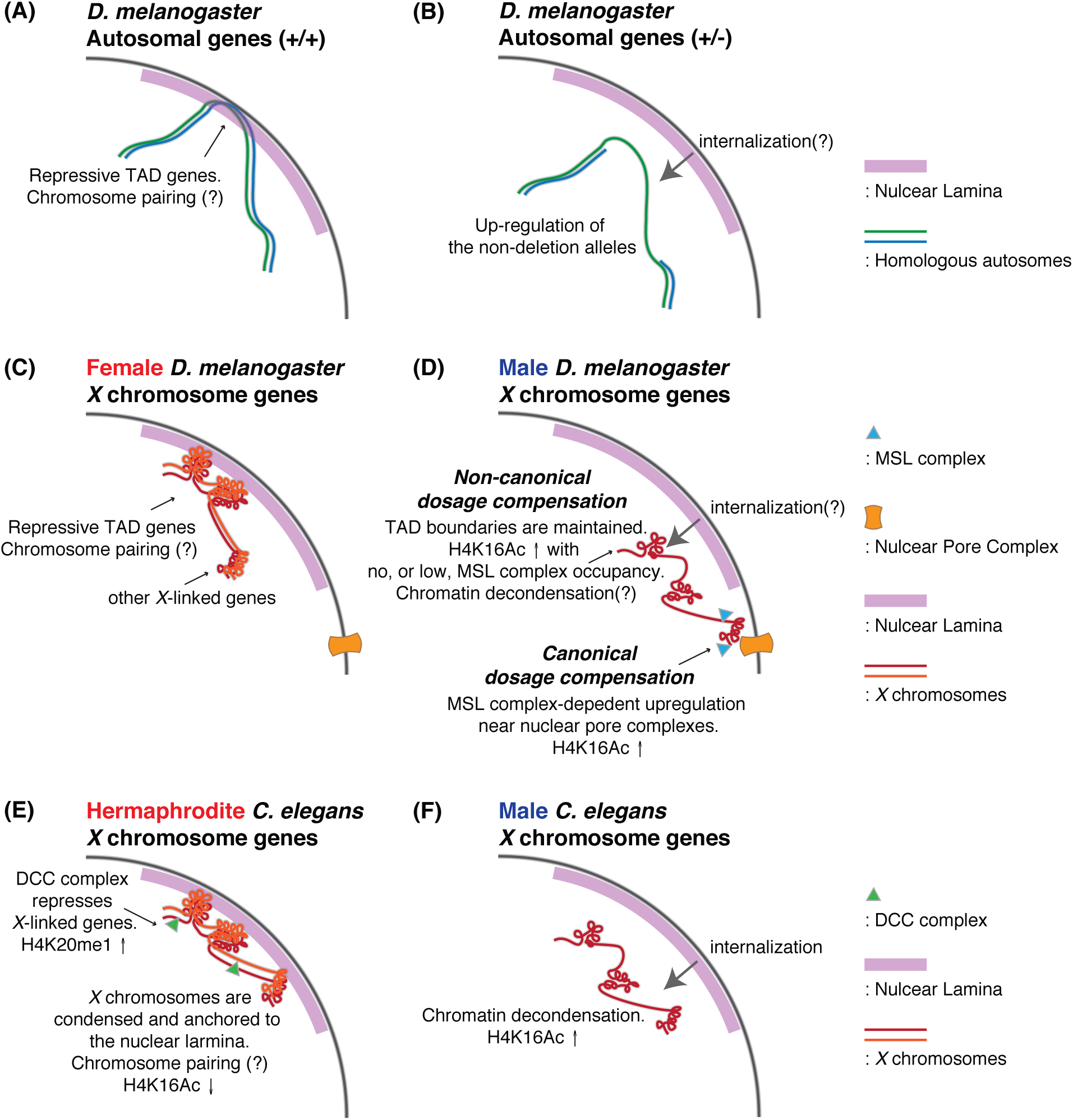
Models demonstrating the parallelism among dosage compensation of autosomal dosage compensation in hemizygous *D. melanogaster*, and *X* chromosome dosage compensation in *C. elegans* and *D. melanogaster*. **(A, B)** A proposal of de-repression mediated compensation of one-dose autosomal genes in hemizygous *D. melanogaster* based on our previous study [28]. **(C, D)** A model of *X* chromosome dosage compensation in D. melanogaster based on the current study as well as other references [5,7,14,39,41,48,79,80]. **(E, F)** A model of *X* chromosome dosage compensation in *C. elegans* based on the references [16,20–23,26,27,81].

De-repression of one-dose genes in males is reminiscent of the *C. elegans* dosage compensation mechanism (**Figure 6E,F**). In *C. elegans, XX* individuals are hermaphrodites and *XO* individuals are males. Both *X* chromosomes in hermaphrodites are subjected to dosage compensation control by repression [3,5,16]. The process involves DCC complex-dependent chromatin remodeling in *XX* hermaphrodites [20–22], that includes enrichment for H4K20me1 and depletion for H4K16Ac. In *XO* worms, the *X* shows decondensation [23]. In addition to the chromatin remodeling, there is local positioning of both *X* chromosomes of hermaphrodites to the LADs at the nuclear periphery which contributes to the repression of *X-*linked gene expression; the loss of this tethering results in de-repression of *X-*linked genes in hermaphrodites [27]. The de-repression of *X-*linked genes in tethering mutants of *cec-4* or *lem-*2, which encode a chromodomain protein or a component of nuclear lamina, respectively, results in a less extreme compensation phenotype than DCC mutants, raising the possibility that tethering to the nuclear lamina is an additional or supplemental mechanism to achieve dosage compensation by repression in *XX* individuals [27]. Thematically, this is identical to the non-canonical model for *Drosophila* dosage compensation that we propose.

*X* chromosome dosage compensation by de-repression appears to rely on a general feature of repressive domains, requiring very little evolutionary innovation. As sex chromosomes evolve from an autosomal pair, the sex chromosome specific to the hetergametic sex, becomes recombinationally silent and accumulates inversions, insertions, and pseudogenes that further disrupt pairing [52–54]. As this process occurs, partial dosage compensation by de-repression would be an immediate response, not requiring the evolution of any specific machinery. Improved dosage compensation can evolve to boost gene expression in *XY* males and by enhancing repression in *XX* females. This could account for some of the commonality between *D. melanogaster* and *C. elegans* despite their divergence ~ 1 billion of years ago [55,56].

The MSL complex does not function specifically on the *X* chromosome in the male germline of *D. melanogaster* [57,58], although they may be dosage compensated [59] (but also see [60]). There is a clear depletion of genes with male biased expression in regions of high MSL occupancy [61], but given that these specific MSL sites do not appear to be used in the male germline, the suggestion that MSL drives these genes to other locations seems spurious. We have shown that the regions without MSL entries sites correspond to the repressive TADs. Thus, we propose that *X-*linked genes with male germline functions are more likely to be in repressive TADs, where they can show increased expression as a result of de-repression. Indeed, in our previous results from gene expression profiling of hemizygote files with autosomal deletions [28], we observed that genes with male-biased expression are de-repressed in females. There has been strong evolutionary pressure to relocate genes with male germline function off the *X* chromosomes [62–64]. Those that remain might use de-repression to achieve high expression even on the single *X*.

## Conclusion

Our results collectively suggest that MSL complex-independent *X* chromosome dosage compensation exists in *Drosophila melanogaster*. We suggest that this non-canonical dosage compensation mechanism involves de-repression of one-dose *X* chromosome genes in males, which are repressed in their two-dose state in females. Our results have an implication for the *X* chromosome dosage compensation mechanism before the evolution of the MSL complex.

## Materials and Methods

### TADs information used in this study

We obtained LAD information from [65], HiC domains from [35], and DamID-based chromatin domains from [34]. All these results were generated based on *Drosophila* reference genome release 5. We used Flybase 5.57 gene model [66] in describing genes within such TADs. We defined genes to belong to TADs only when both boundaries of a gene locate in a TAD region. We performed our gene ontology analysis in FlyMine version 45.1 [67]. Results in the Additional File 1 represents significantly enriched terms, adjusted *p* value < 0.05, after Holm-Bonferroni correction. Hi-C based TAD boundary information for *S2* and *Kc* cells were obtained from [47].

### *Drosophila* cell line data from modENCODE studies

We used our previous results on RNA-Seq expression profiles of *Drosophila Kc* and *S2* cells [37] for this study after updating gene IDs to Flybase 5.57. We used FPKM > 1 as an expression cutoff based on the top 99th percentile of the intergenic FPKM signals (0.87 and 0.98 for *Kc* and *S2* cells, respectively). We used following chromatin immunoprecipitation (ChIP)-on-chip results from modENCODE study (model organism ENcyclopedia of DNA Elements, [40]. modENCODE submission IDs 3043 and 3044 for MOF binding in *Kc* and *S2* cells, respectively, ID 318 for Histone H4K16 acetylation in *Kc* cells, and IDs 319 and 320 for *S2* cells. In our description of H4K16 acetylation levels in *S2* cells in **Figure 2**, we used median values from these two different submissions. We obtained MSL-1 binding results from modENCODE submission ID 3293. These datasets can also be obtained from Gene Expression Omnibus (GEO, [68] with these accession IDs: GSE27805-6, GSE20797-9, and GSE32762. modENCODE study [40] provided smoothed log-intensity values between ChIP signal and the input signal, called M values, whose processed mean is shifted to 0. We used median M values within gene boundaries in describing MOF/MSL-1 binding or H4K16 acetylation in **Figure 2A-I**. MOF binding and H4K16 acetylation enriched/not-enriched regions in **Figure 3** directly followed peak calls from the original study.

### Salivary gland expression profiles and ChIP-Seq results

We obtained microarray expression profiling and ChIP-Seq results from the 3rd instar larva salivary glands for MOF binding and Histone H4K16 acetylation from [36]. The gene expression profiles were provided as GCRMA (GC Robust Multi-array Average, [69]-normalized signal intensities, and we used the top 95 percentiles of signals from *non-Drosophila* control probes as an expression cutoff. We demonstrated the median values from three replicates in **Figure 1C-E**. The original results can be found from ArrayExpress [70] with accession ID of E-MEXP-3506. ChIP-Seq results for MOF binding and H4K16 acetylation, from the same study, can be accessed with ArrayExpress ID E-MTAB-911. In the result, the authors performed analysis with DESeq [71] to calculate log2 fold changes between ChIP and input samples for non-overlapping 25 bp windows across the genome. We used median values of such log2 fold changes within gene boundaries in describing the ChIP results in **Figure 2J-O**.

### MSL entry sites

We used 150 CES that were characterized by ChIP-chip and ChIP-Seq studies [14] to generate a position weight matrix for DCC binding using MEME (Multiple EM for Motif Elicitation) suite version 4.11.2 [72]. We set the length of the motif to be 21 bp to match with the original CES study. Using the position weight matrix, we identified locations with MREs across the *Drosophila* genome release 5. We used FIMO 4.11.2 (Find Individual Motif Occurrences, [73] in this identification with *Expect* value (E-value) threshold of 1.0e-05. In our description of MRE/CES occurrence in **Figure 2**, we randomly shuffled positions of TADs on *X* chromosome genome using Bedtools 2.26.0 [74] while preserved the sizes of TADs. The results in **Figure 2R,S** demonstrate overlap between such shuffled TADs and MRE/CES from 2,000 randomizations.

### *S2* cell RNAi results for MSL knockdown

We used *mof, msl-1, msl-3* knockdown results from a microarray study [36], ArrayExpress E-MEXP-1505). For the estimation of gene expression changes, we used Robust Multi-array Average (RMA, [75] method for background adjustment and normalization, and filtered out genes of which FPKM value is less than 1 from the *S2* cell RNA-Seq result [37]. We use R limma package version 3.28.21 [76] as in the official manual for our differential expression analysis. We obtained the microarray study of the *msl-2* knockdown data from [41]. We conducted same data handling process as above. We also re-analyzed RNA-Seq results from [42] (GEO GSE16344). We used HISAT 2.0.4 [77] for the mapping of sequencing reads to *Drosophila* genome release 5. We used a parameter for unpaired sequencing (-U) in running HISAT. We measured gene-level read abundances with HTSeq 0.6.1 [77] with the default setting. From the counting result, we used polyA+ protein coding genes that have more than 1 count per million mapped reads from any of the four samples (two controls and two RNAi) in our differential expression analysis. We performed differential expression analysis using DESeq2 [78]. In **Figure 3** and **Figure 5**, we demonstrated genes of which expression is more than 1 FPKM, which we also used to filter microarray results from MSL knockdown.

### List of abbreviations

CES:Chromosome Entry Sites, ChIP: Chromatin Immunoprecipitation, DCC: Dosage Compensation Complex, FPKM: Fragments Per Kilobase of transcript per Million mapped reads, GEO: Gene Expression Omnibus, GO: Gene Ontology, H4K16Ac: Histone H4 Lysine 16 Acetylation, HAS: High-Affinity Sites, LAD: Lamina-Associated Domains, MLE: Maleless, modENCODE: model organism ENcyclopedia of DNA Elements, MOF: Males absent on the first, MRE: MSL-recognition element, MSL: Male specific lethal complex, NAR: Nucleoporin-Associated Region, NSL: Non-Specific Lethal, TAD: Topologically Associated Domain.

## Declarations

### Ethics approval and consent to participate

Not applicable

### Consent for publication

Not applicable

### Availability of data and material

The datasets analysed during the current study are available in the GEO and ArrayExpress repositories. We used modENCODE ChIP-chip results that are available in GEO with these accession IDS: GSE27805-6, GSE20797-9, and GSE32762. The salivary glands results are in ArrayExpress (E-MTAB-911 and E-MEXP-3506). We re-analyzed MSL complex knockdown results from GEO GSE16344 and ArrayExpress E-MEXP-1505.

### Competing interests

The authors declare that they have no competing interests

## Funding

This work was supported by the Intramural Research Programs of the National Institutes of Health (NIH), National Institute of Diabetes and Digestive and Kidney Diseases, to BO, and Korean Visiting Scientist Training Award (KVSTA, HI13C1282) to HL.

## Authors’ contributions

HL and BO conceived of the idea and designed the analyses. HL performed computational analysis on the presented results. HL and BO interpreted and wrote the manuscript.

## Acknowledgements

We thank the members of the Oliver lab for their helpful discussions and Dr. Per Stenberg and Dr. Sergey V. Razin for kindly sharing processed results from their studies. We utilized the high-performance computational capabilities of the Biowulf linux cluster at the NIH, Bethesda, MD. This research was supported by the Intramural Research Program of the NIH, The National Institute of Diabetes and Digestive and Kidney Diseases (NIDDK).

